# Chemoproteomic Profiling Reveals that Anti-Cancer Natural Product Dankastatin B Covalently Targets Mitochondrial VDAC3

**DOI:** 10.1101/2023.02.11.528139

**Authors:** Bridget P. Belcher, Paulo A. Machicao, Binqi Tong, Emily Ho, Julia Friedli, Brian So, Helen Bui, Yosuke Isobe, Thomas J. Maimone, Daniel K. Nomura

**Affiliations:** Department of Chemistry, University of California, Berkeley, Berkeley, CA 94720 USA; Novartis-Berkeley Translational Chemical Biology Institute, Berkeley, CA 94704 USA; Innovative Genomics Institute, Berkeley, CA 94704 USA; Department of Molecular and Cell Biology, University of California, Berkeley, Berkeley, CA 94720 USA

**Keywords:** chemoproteomics, activity-based protein profiling, natural products, gymnastatins, dankastatin B, VDAC, RNF114

## Abstract

Chlorinated gymnastatin and dankastatin alkaloids derived from the fungal strain *Gymnascella dankaliensis* have been reported to possess significant anti-cancer activity but their mode of action is unknown. These members possess electrophilic functional groups that may undergo covalent bond formation with specific proteins to exert their biological activity. To better understand the mechanism of action of this class of natural products, we mapped the proteome-wide cysteine-reactivity of the most potent of these alkaloids, dankastatin B, using activity-based protein profiling chemoproteomic approaches. We identified a primary target of dankastatin B in breast cancer cells as cysteine C65 of the voltage-dependent anion selective channel on the outer mitochondrial membrane VDAC3. We demonstrated direct and covalent interaction of dankastatin B with VDAC3. VDAC3 knockdown conferred hyper-sensitivity to dankastatin B-mediated anti-proliferative effects in breast cancer cells indicating that VDAC3 was at least partially involved in the anti-cancer effects of this natural product. Our study reveals a potential mode of action of dankastatin B through covalent targeting of VDAC3 and highlight the utility of chemoproteomic approaches in gaining mechanistic understanding of electrophilic natural products.

---

Natural products have served as a rich source of medicinal agents for various diseases including cancer. Among these natural products are those that possess electrophilic functional groups that can covalently act with nucleophilic amino acids within proteins to exert their biological activity. These covalently-acting natural products can often selectively engage with unique ligandable sites within proteins and provide sustained target engagement. Members displaying this reactivity include β-lactam antibiotics such as penicillin that covalently target transpeptidases, epoxomicin that irreversibly inhibits the proteasome, fumagillins that target methionine aminopeptidase 2, Wortmannin that targets kinases including phosphoinositide 3-kinase, and acetylsalicyclic acid that acetylates a serine within cyclooxygenases ^1,2^. Despite the identification of numerous natural products possessing both protein-reactive functional groups and therapeutically relevant biological activities, the biological targets and mechanism(s) of action of most covalently-acting natural products are unknown.

Among these natural products, we became interested in chlorinated gymnastatin and dankastatin alkaloids derived from the fungus *Gymnascella dankaliensis* isolated from the sponge *Halichondria japonica* ^3–6^. Many of these tyrosine-derived alkaloids have been reported to possess significant anti-cancer activity, but their mechanisms of action remain unknown ^3–6^. These gymnastatin and dankastatin alkaloids possess many distinct electrophilic functional groups, including chloroenone, α-chloroketone, epoxyketone, lactol, acetal, and α , β , γ , δ-unsaturated amide moieties that can potentially react with nucleophilic amino acids such as cysteines (See Figure 1). We recently reported total syntheses of many of these alkaloids^7^, and as such, we delved into the anti-cancer and mechanistic characterization of these compounds. Herein we provide the first proteomic profiling and target annotation for a member of this class of cytotoxic natural products.

**Figure 1.**
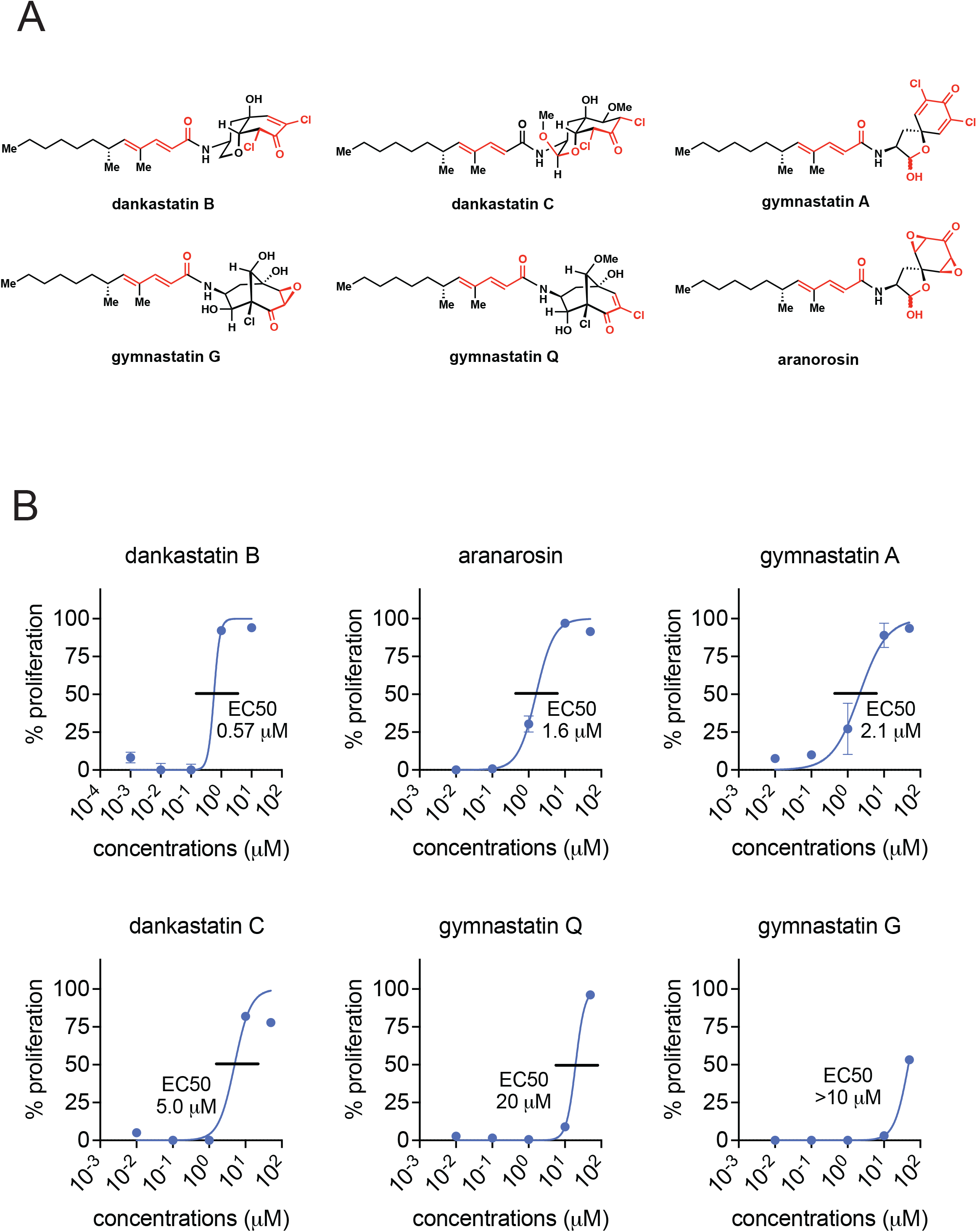
Anti-proliferative effects of gymnastatin and dankastatin alkaloids in breast cancer cells. **(A)** Structures of gymnastatin and dankastatin alkaloids tested in this study. Shown in red are potential electrophilic sites on the molecules. **(B)** Inhibition of MDA-MB-231 cell proliferation. MDA-MB-231 breast cancer cells were treated with DMSO vehicle or natural products for 24 h and cell proliferation was assessed by Hoechst staining. 50 % effective concentration (EC50) are shown. Data shown are average ± sem from n=6 biological replicates/group.

We first tested the anti-proliferative activity of five gymnastatin and dankastatin alkaloids—dankastatin B, gymnastatin A, dankastatin C, gymnastatin Q, and gymnastatin G–as well as the related natural product aranorosin, in triple-negative breast cancer cells **(Figure 1A-1B)**. All of these compounds showed dose-responsive impairment in MDA-MB-231 breast cancer cell proliferation with varying degrees of sensitivity with dankastatin B showing the greatest potency with a 50 % effective concentration (EC50) of 0.57 μM and gymnastatin G showing the weakest potency with EC50 >10 μM **(Figure 1B)**. Given the structural similarity between these classes of natural products, we were intrigued by differences in anti-proliferative effects and the potential for differences in their potential protein target profiles. We focused on mapping the proteome-wide cysteine-reactivity of the most potent dankastatin B and the least potent gymnastatin G using activity-based protein profiling (ABPP) chemoproteomic approaches, namely isotopic tandem orthogonal proteolysis-ABPP (isoTOP-ABPP) ^7–9^. Out of >2000 cysteines profiled and quantified across each experiment, dankastatin B (treated at 10 μM) revealed only one target—cysteine 65 on the voltage-dependent anion-selective channel protein 3 (VDAC3)—with a control-to-treated ratio >4, indicating >75 % target engagement, with statistical significance (adjusted p-value <0.05) **(Figure 2, Table S1).** We note that there were an additional 18 targets that showed a ratio >2 with adjusted p-value <0.05, indicating that there are likely additional targets beyond VDAC3 of dankastatin B in breast cancer cells that show ∼50-75 % target engagement **(Table S1)**. Interestingly, among these targets was also C76 of VDAC2, another isoform of VDAC3. VDACs are cysteine-rich, pore-forming proteins that coat the outer membrane of the mitochondria and are major regulators for metabolite exchange between the cytosol and mitochondria and previous studies have shown that VDAC silencing lead to impaired mitochondrial function, including ATP production, calcium flux, reactive oxygen stress balance and apoptosis ^10^. In contrast, gymnastatin G did not reveal any targets with ratios >4 and only two targets with ratios >2 comparing cells treated with vehicle versus 50 μM of gymnastatin G **(Figure S1, Table S2)**. These two targets with ratios >2 was C70 of RPL27A and C20 of TUB1A—both unrelated to VDAC2/3. The ratio observed for VDAC3 C65 and VDAC2 C70 with gymnastatin G treatment was 0.29 and 0.61, respectively, and not statistically significant. These data may suggest that gymnastatin G is either less reactive, does not possess potency against dankastatin B targets, or may be unstable in living cells, and may potentially explain why dankastatin B showed more potent anti-proliferative effects compared to gymnastatin G.

**Figure 2.**
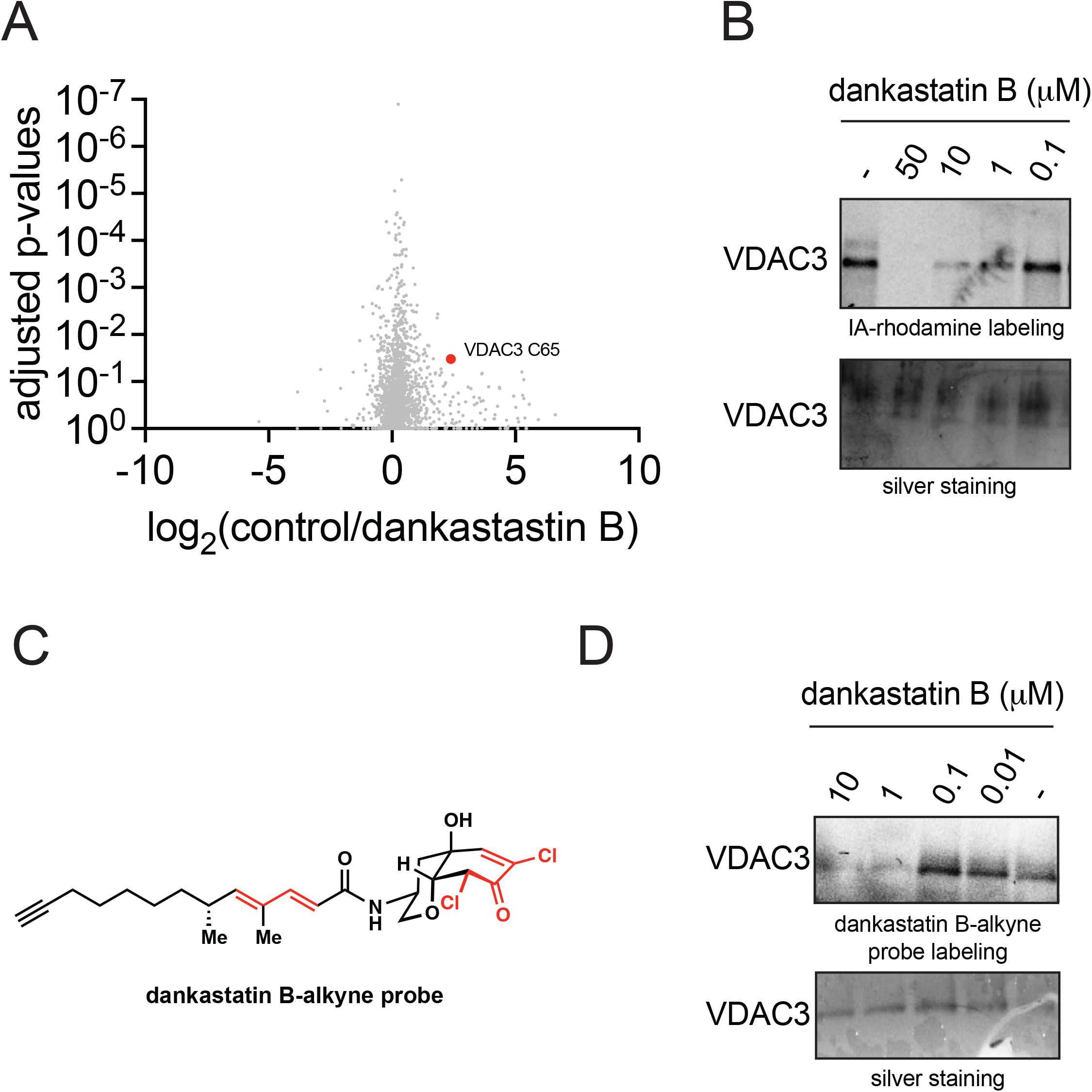
Chemoproteomic profiling reveals VDAC3 as a target of dankastatin B. **(A)** isoTOP-ABPP analysis of dankastatin B in MDA-MB-231 breast cancer cells. MDA-MB-231 breast cancer cells were treated with DMSO vehicle or dankastatin B (10 μM) for 1 h. Resulting cell lysates were treated with the cysteine-reactive alkyne-functionalized iodoacetamide probe (IA-alkyne) and subsequently taken through the isoTOP-ABPP procedure and LC-MS/MS analysis. Control (isotopically light) versus treated (isotopically heavy) probe-modified peptide ratios and adjusted p-values were calculated and shown. Shown is red are the targets that showed control/treated ratio >4 with adjusted p-values <0.05 from n=6 biological replicates/group. Only C65 of VDAC3 fit these criteria. **(B)** Gel-based ABPP analysis of dankastatin B against pure VDAC3 protein. Pure VDAC3 protein was pre-incubated with DMSO or dankastatin B for 30 min prior to labeling of protein with a rhodamine-functionalized iodoacetamide (IA-rhodamine) probe (1 μM) for 1 h. Proteins were resolved on SDS/PAGE and subsequently visualized by in-gel fluorescence and protein loading was assessed by silver staining. **(C)** Shown is a structure of the alkyne-functionalized dankastatin B probe, dankastatin B-alkyne. Shown in red are the potential electrophilic sites. **(D)** Dankastatin B-alkyne probe labeling of VDAC3 and competition of probe labeling by dankastatin B. Pure VDAC3 protein was pre-incubated with DMSO or dankastatin B for 30 min prior to incubation with dankastatin B-alkyne (50 μM) for 30 min. An azide-functionalized rhodamine was subsequently appended onto probe-labeled proteins by copper-catalyzed azide-alkyne cycloaddition, proteins were resolved by SDS/PAGE, and visualized by in-gel fluorescence. Protein loading was assessed by silver staining. Shown in **(B, D)** are representative gels from n=3 biological replicates/group.

Intrigued by these results, we next biochemically confirmed the interaction of dankastatin B with VDAC3 and VDAC2. First, we showed dose-responsive competition of dankastatin B against the binding of a rhodamine-functionalized cysteine-reactive iodoacetamide probe to pure VDAC3 and VDAC2 by gel-based ABPP **(Figure 2B; Figure S2A).** Second, we synthesized an alkyne-functionalized probe of dankastatin B **(Figure 2C)** and showed that this probe directly and covalently binds to pure VDAC3 and VDAC2 protein and that this labeling is also dose-responsively competed out by dankastatin B **(Figure 2D; Figure S2B-S2C)**.

While we do not think that VDAC3 is the only target responsible for the anti-proliferative effects of dankastatin B, we nonetheless sought to determine whether VDAC3 plays at least a partial role in the anti-cancer outcomes of this natural product. We showed that VDAC3 knockdown by short-hairpin RNA interference led to hyper-sensitivity to dankastatin B effects in breast cancer cells **(Figure 3A-3B).** We do not know currently whether dankastatin B, through targeting C65, inhibits or activates VDAC3 channel activity, its effects on pore assembly with the other VDAC proteins, or its effects on mitochondrial membrane potential or permeability. However, given the various roles of VDAC3 in modulating mitochondrial function and apoptosis ^10^, we conjectured that covalent targeting of VDAC3 may cause mitochondria-dependent apoptosis of breast cancer cells. Mitochondrial apoptosis occurs through the intrinsic apoptotic pathway that involves activation of caspase 9 ^11^. Consistent with dankastatin B causing mitochondria-dependent intrinsic apoptosis, we observed significant rescue of dankastatin B anti-proliferative effects with a caspase 9-selective inhibitor z-LEHD-FMK **(Figure 3C).**

**Figure 3.**
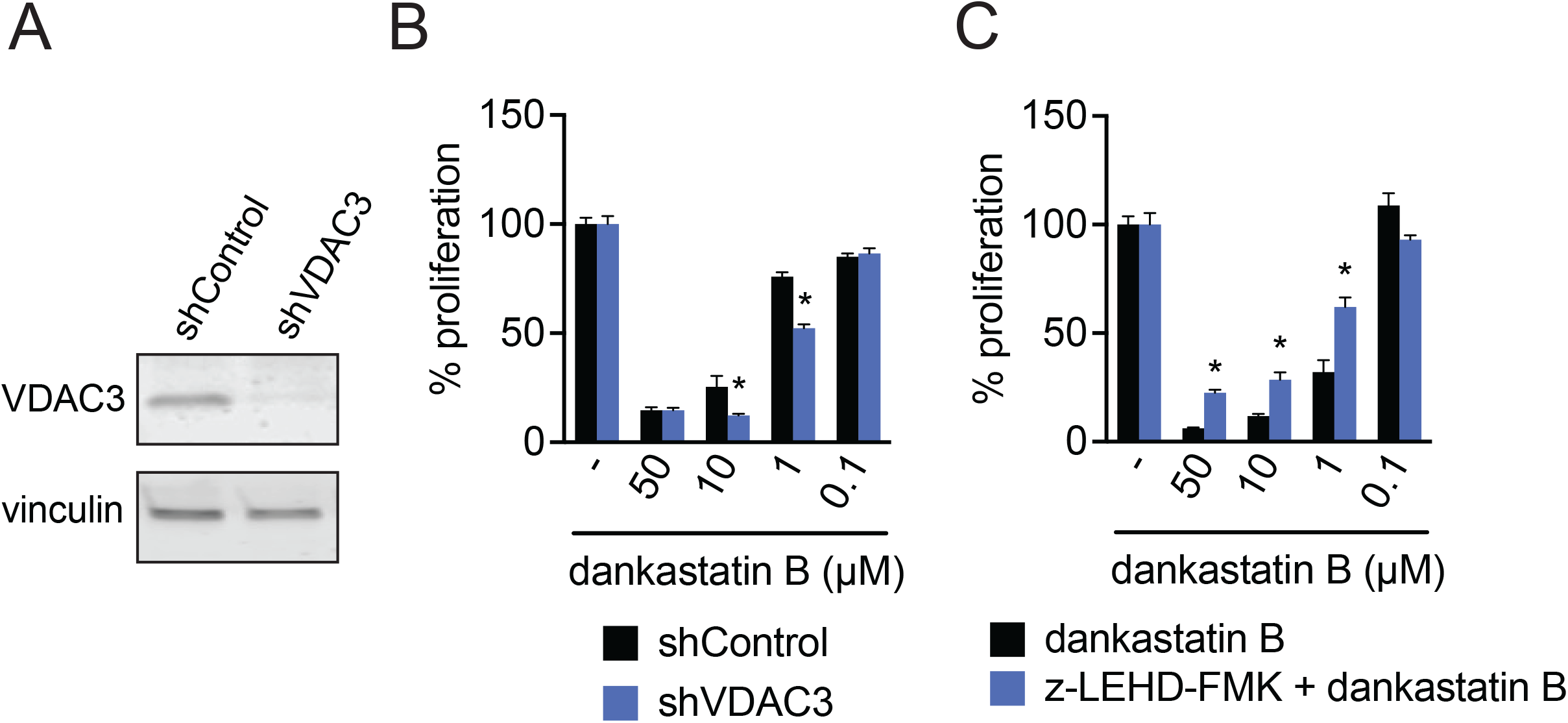
Role of VDAC3 in anti-proliferative effects. **(A)** Knockdown of VDAC3. VDAC3 was stably knocked down by short hairpin RNA (shRNA). Knockdown of VDAC3 was validated by Western blotting compared to a loading control vinculin. **(B)** MDA-MB-231 cell proliferation in shControl versus shVDAC3 cells. MDA-MB-231 shControl and shVDAC3 cells were treated with DMSO vehicle or dankastatin B for 8 h and cell proliferation was assessed by Hoechst stain. **(C)** MDA-MB-231 cell proliferation with caspase 9 inhibitor z-LEHD-FMK. MDA-MB-231 cells were pre-treated with DMSO vehicle or z-LEHD-FMK for 2 h prior to treatment with DMSO vehicle or dankastatin B for 8 h. Cell proliferation was read out by Hoechst stain. Data shown are average ± sem from n=6 biological replicates/group. Significance is shown as *p<0.05 compared to the equivalent dankastatin B concentration for shControl in **(B)** or compared to treatment with each dankastatin B concentration alone in **(C)**.

Overall, our study investigated potential mode of action of the electrophilic natural product dankastatin B, a family member of an interesting class of chlorinated gymnastatin and dankastatin alkaloids, wherein we showed relatively selective covalent targeting of C65 on the mitochondrial pore-forming VDAC3. We also demonstrate direct covalent binding to a VDAC3 isoform VDAC2 on C76. We also demonstrated that VDAC3 was at least partially involved in the anti-proliferative effects observed with dankastatin B. Interestingly, we showed that a closely related natural product gymnastatin G that demonstrated poorer anti-cancer potency, showed very little reactivity compared to dankastatin B. Of future interest is to further investigate the structure-activity relationships of gymnastatin and dankastatin B alkaloids and their effects on VDAC2/3. Also worthy of further investigation is whether targeting key cysteines within VDAC proteins may represent a potential therapeutic strategy for cancer. Our study also further highlights the utility of chemoproteomic platforms to discover the mode of action of biologically active natural products.

## Acknowledgement

We thank the members of the Nomura Research Group and Novartis Institutes for BioMedical Research for critical reading of the manuscript. This work was supported by Novartis Institutes for BioMedical Research and the Novartis-Berkeley Translational Chemical Biology Institute (NB-TCBI) for all listed authors. This work was also supported by the Nomura Research Group and the Mark Foundation for Cancer Research ASPIRE Award for DKN, BB. We acknowledge support from grants from the National Institutes of Health (R01GM136945 for DKN and TJM and an NIGMS diversity supplement to PAM as well as grant R35CA263814 for DKN). We also thank Drs. Hasan Celik, Alicia Lund, and UC Berkeley’s NMR facility in the College of Chemistry (CoC-NMR) for spectroscopic assistance. Instruments in the College of Chemistry NMR facility are supported in part by NIH S10OD024998.

## Author Contributions

BPB, TJM, DKN conceived of the project idea, designed experiments, performed experiments, analyzed and interpreted the data, and wrote the paper. BPB, PAM, BT, EH, JF, BS, HB, YI, DKN performed experiments, analyzed and interpreted data, and provided intellectual contributions.

## Methods

### Cell Culture

MDA-MB-231 cells were obtained from the UC Berkeley Cell Culture Facility and were cultured in L15 medium (HyClone) containing 10% (v/v) fetal bovine serum (FBS) and maintained at 37°C with 0% CO_2_. HEK293T cells were obtained from the UC Berkeley Cell Culture Facility and were cultured in Dulbecco’s Modified Eagle Medium (DMEM) containing 10% (v/v) FBS and maintained at 37°C with 5% CO_2_.

### Proliferation Assays

Cell proliferation assays were performed using Hoechst 33342 dye (Invitrogen) according to the manufacturer’s protocol and as previously described^12^. MDA-MB-231 cells were seeded into 96-well plates (20,000 cells/well) in a volume of 150 μL and allowed to adhere overnight. Cells were treated with an additional 50 μL of media containing DMSO vehicle or a 1:250 dilution of 1,000x alkaloid natural product stock. After the incubation period, the media was removed from each well and 100 μL of staining solution containing 10% formalin and Hoechst 33342 dye was added to each well. The cells were incubated for 15 min in the dark at room temperature, the staining solution was removed, and the fixed cells were washed with PBS before imaging. Fluorescence was read with a Tecan Spark plate reader (λex = 350 nm, λem = 461 nm).

### Assessing Apoptosis in Breast Cancer Cells

MDA-MB-231 cells were seeded into 96-well plates (20,000 cells/well) in a volume of 150 μL and allowed to adhere overnight. Cells were pre-treated with caspase 9 inhibitor z-LEHD-FMK (20 μM final concentration, Abcam) for 2 h. Cells were then treated with dankastatin B (1,000x stock in DMSO) or DMSO vehicle. After an 8 h incubation, the media was removed from each well and 100 μL of staining solution containing 10% formalin and Hoechst 33342 dye was added to each well. The cells were incubated for 15 min in the dark at room temperature, the staining solution was removed, and the fixed cells were washed with PBS before imaging. Fluorescence was read with a Tecan Spark plate reader (λex = 350 nm, λem = 461 nm).

### Preparation of Cell Lysates

Cells were washed with cold PBS, scraped, and pelleted by centrifugation (1400 g, 5 min, 4°C). Pellets were resuspended in cold PBS, lysed by sonication or using RIPA buffer, and clarified by centrifugation (12,000 g, 10 min, 4°C). Lysate was transferred to low-adhesion microcentrifuge tubes. Proteome concentrations were determined using BCA assay and lysate was diluted with PBS to appropriate working concentrations.

### Western Blotting

Antibodies to VDAC3 (Invitrogen, PA5-51156), VDAC2 (Abcam, Ab1216120), and vinculin (Bio-Rad, MCA465GA) were obtained from various commercial sources and dilutions were prepared per the manufacturers’ recommendations. Proteins were resolved by SDS/PAGE and transferred to nitrocellulose membranes using the Trans-Blot Turbo transfer system (Bio-Rad). Membranes were blocked in 5% BSA in Tris-buffered saline containing Tween 20 (TBS-T) solution for 30 min at RT, washed in TBS-T, and probed with primary antibody for 1.5 h at RT. Membranes were washed again with TBS-T and incubated for 1 h at RT in the dark with IR680- and IR800-conjugated secondary antibodies at 1:10,000 dilution in 5% BSA in TBS-T. Following 3 additional TBS-T washes, bots were visualized using an Odyssey Li-Cor fluorescent scanner.

### isoTOP-ABPP Chemoproteomics

IsoTOP-ABPP studies were performed as previously reported^7,9,12^. Cells were lysed by probe sonication in PBS and protein concentrations were measured by BCA assay. Cells were treated for 1 h with either DMSO vehicle or alkaloid natural product before cell collection and lysis. Proteomes were subsequently labeled with IA-alkyne labeling (100 μM) for 1 h at RT. CuAAC was used by sequential addition of tris(2-carboxyethyl)phosphine (1 mM, Strem), tris[(1-benzyl-1H-1,2,3-triazol-4-yl)methyl]amine (34 μM, Sigma), copper (II) sulfate (1 mM, Sigma) and biotin-linker-azide – the linker functionalized with a tobacco etch virus (TEV) protease recognition sequence as well as an isotopically light or heavy valine for treatment of control or treated proteome, respectively. After CuAAC, proteomes were precipitated by centrifugation at 6,500 g, washed in ice-cold methanol, combined in a 1:1 control/treated ratio, washed again, then denatured and resolubilized by heating in 1.2% SDS/PBS to 90°C for 5 min. Insoluble components were precipitated by centrifugation at 6,500 g and soluble proteome was diluted in 5 mL 0.2% SDS/PBS. Labeled proteins were bound to streptavidin-agarose beads (170 μL resuspended beads per sample, Thermo Fisher) while rotating overnight at 4°C. Bead-linked proteins were enriched by washing three times each in PBS and water, then resuspended in 6 M urea/PBS (Sigma), reduced in DTT (9.26 mM, Thermo Fisher), and alkylated with iodoacetamide (18 mM, Sigma) before being washed and resuspended in 2 M urea/PBS and trypsinized overnight with 0.5 μg/μL sequencing-grade trypsin (Promega). Tryptic peptides were eluted. Beads were washed three times each in PBS and water, washed in TEV buffer solution (water, TEV buffer, 100 μM dithiothreitol) and resuspended in buffer with Ac-TEV protease (Invitrogen) and incubated overnight. Peptides were diluted in water, acidified with formic acid (1.2 M, Fisher), and prepared for analysis.

### isoTOP-ABPP Mass Spectrometry Analysis

Peptides from all chemoproteomic experiments were pressure-loaded onto a 250 μm inner diameter fused silica capillary tubing packed with 4 cm of Aqua C18 reverse-phase resin (Phenomenex, 04A-4299), which had been previously equilibrated on an Agilent 600 series high-performance liquid chromatograph using the gradient from 100% buffer A to 100% buffer B over 10 min, followed by a 5 min wash with 100% buffer B and a 5 min wash with 100% buffer A. The samples were then attached using a MicroTee PEEK 360 μm fitting (Thermo Fisher Scientific, p-888) to a 13 cm laser-pulled column packed with 10 cm Aqua C18 reverse-phase resin and 3 cm of strong-cation exchange resin for isoTOP-ABPP studies. Samples were analyzed using an Q Exactive Plus mass spectrometer (Thermo Fisher Scientific) using a five-step Multidimensional Protein Identification Technology (MudPIT) program, using 0, 25, 50, 80 and 100% salt bumps of 500 mM aqueous ammonium acetate and using a gradient of 5–55% buffer B in buffer A (buffer A: 95:5 water/acetonitrile, 0.1% formic acid; buffer B 80:20 acetonitrile/water, 0.1% formic acid). Data were collected in data-dependent acquisition mode with dynamic exclusion enabled (60 s). One full mass spectrometry (MS1) scan (400–1,800 mass-to-charge ratio (*m/z*)) was followed by 15 MS2 scans of the *n*th most abundant ions. Heated capillary temperature was set to 200 °C and the nanospray voltage was set to 2.75 kV.

Data were extracted in the form of MS1 and MS2 files using Raw Extractor v.1.9.9.2 (Scripps Research Institute) and searched against the Uniprot human database using ProLuCID search methodology in IP2 v.3-v.5 (Integrated Proteomics Applications, Inc.)^13^. Cysteine residues were searched with a static modification for carboxyaminomethylation (+57.02146) and up to two differential modifications for methionine oxidation and either the light or heavy TEV tags (+464.28596 or +470.29977, respectively). Peptides were required to be fully tryptic peptides and to contain the TEV modification. ProLUCID data were filtered through DTASelect to achieve a peptide false-positive rate below 5%. Only those probe-modified peptides that were evident across two out of three biological replicates were interpreted for their isotopic light to heavy ratios. For those probe-modified peptides that showed ratios greater than two, we only interpreted those targets that were present across all three biological replicates, were statistically significant and showed good quality MS1 peak shapes across all biological replicates. Light versus heavy isotopic probe-modified peptide ratios are calculated by taking the mean of the ratios of each replicate paired light versus heavy precursor abundance for all peptide-spectral matches associated with a peptide. The paired abundances were also used to calculate a paired sample *t*-test *P* value in an effort to estimate constancy in paired abundances and significance in change between treatment and control. *P* values were corrected using the Benjamini–Hochberg method.

### Gel-Based ABPP

Gel-Based ABPP methods were performed as previously described. Recombinant pure human VDAC2 was obtained from NOVUS (NBP1-89477PEP) and recombinant pure human VDAC3 was obtained from Abnova (H00007419-P01). Proteins (0.1 μg) were pre-treated with either DMSO vehicle or dankastatin B for 30 min at 37°C in an incubation volume of 25 μL PBS, and subsequently treated with IA-Rhodamine (1 μM, Setareh Biotech) for 1 h in the dark at room temperature. The reactions were stopped by the addition of 4x reducing Laemmli SDS sample loading buffer (Alfa Aesar) and heated at 95°C for 5 min. The samples were separated on precast 4-20% Criterion TGX gels (Bio-Rad). Probe-labeled proteins were analyzed by in-gel fluorescence using a ChemiDoc MP (Bio-Rad). Protein loading was assessed by silver stain.

For competitive gel-based ABPP experiments, proteins (0.1 μg) were pre-treated with either DMSO vehicle or dankastatin B for 30 min at 37°C in an incubation volume of 25 μL PBS, and subsequently treated with an appropriate concentration of dankastatin B-alkyne for 30 min at 37°C. CuAAC was performed to append rhodamine-azide (1 μM final concentration) onto alkyne probe-labeled proteins. Samples were then diluted with 4x reducing Laemmli SDS sample loading buffer and heated at 95°C for 5 min. The samples were separated on precast 4-20% Criterion TGX gels (Bio-Rad). Prior to analysis, gels were fixed in a solution of 10% acetic acid and 30% ethanol for 1 h. Probe-labeled proteins were analyzed by in-gel fluorescence using a ChemiDoc MP (Bio-Rad). Protein loading was assessed by silver stain.

### Knockdown Studies

Short-hairpin oligonucleotides were used to knock down the expression of VDAC2 and VDAC3 in MDA-MB-231 cells. For lentivirus production, lentiviral plasmids and packaging plasmids (pMD2.5G, Addgene catalog no. 12259 and psPAX2, Addgene catalog no. 12260) were transfected into HEK293T cells using Lipofectamine 2000 (Invitrogen). Lentivirus was collected from filtered cultured medium and used to infect the target cell line with 1:1000 dilution of polybrene. Target cells were selected over 2 d with 1 μg/ml of puromycin. The short-hairpin sequences which were used for generation of the knockdown lines were:

VDAC2: GCTACCCACCAATAATGAAAT (Sigma VDAC2 MISSION shRNA Bacterial Glycerol Stock, TRCN0000151244).

VDAC3: CAGGCAACCTAGAAACCAAAT (Sigma VDAC3 MISSION shRNA Bacterial Glycerol Stock, TRCN0000141213).

MISSION TRC1.5 pLKO.1-puro Non-Mammalian shRNA Control (Sigma) was used as a control shRNA.

