## Supporting Information for "Chemoproteomic Profiling Reveals that Anti-Cancer Natural Product Dankastatin B Covalently Targets Mitochondrial VDAC3"

**Supporting Tables**

**Table S1. Chemoproteomic profiling of dankastatin B.** MDA-MB-231 breast cancer cells were treated with DMSO vehicle or dankastatin B (10 μM) for 2 h. Resulting cell lysates were treated with the cysteine-reactive alkyne-functionalized iodoacetamide probe (IA-alkyne) and subsequently taken through the isoTOP-ABPP procedure and LC-MS/MS analysis. Data are from n=6 biological replicates. Tab 1 shows all probe-modified peptides detected and quantified. Tab 2 and Tab 3 both show probe-modified peptides that were evident in at least 3 out of 6 biological replicates. Tab 2 highlighted proteins/sites show ratio >2 with adjusted p-value <0.05. Tab 3 highlighted proteins/sites show ratio >4 with adjusted p-value <0.05.

**Table S2. Chemoproteomic profiling of gymnastatin G.** MDA-MB-231 breast cancer cells were treated with DMSO vehicle or gymnastatin G (50 μM) for 3 h. Resulting cell lysates were treated with the cysteine-reactive alkyne-functionalized iodoacetamide probe (IA-alkyne) and subsequently taken through the isoTOP-ABPP procedure and LC-MS/MS analysis. Data are from n=3 biological replicates. Tab 1 shows all probe-modified peptides detected and quantified. Tab 2 and Tab 3 both show probe-modified peptides that were evident in at least 2 out of 3 biological replicates. Tab 2 highlighted proteins/sites show ratio >2 with adjusted p-value <0.05. Tab 3 highlighted proteins/sites show ratio >4 with adjusted p-value <0.05.

**Supporting Figures**

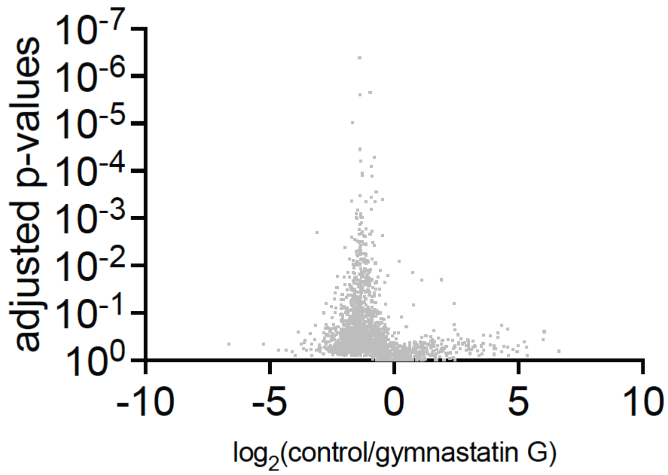


**Figure S1. Chemoproteomic profiling of gymnastatin G.** MDA-MB-231 breast cancer cells were treated with DMSO vehicle or gymnastatin G (50 μM) for 3 h. Resulting cell lysates were treated with the cysteine-reactive alkyne-functionalized iodoacetamide probe (IA-alkyne) and subsequently taken through the isoTOP-ABPP procedure and LC-MS/MS analysis. Data are from n=3 biological replicates. Tab 1 shows all probe-modified peptides detected and quantified. Tab 2 and Tab 3 both show probe-modified peptides that were evident in at least 2 out of 3 biological replicates. Tab 2 highlighted proteins/sites show ratio >2 with adjusted p-value <0.05. Tab 3 highlighted proteins/sites show ratio >4 with adjusted p-value <0.05.

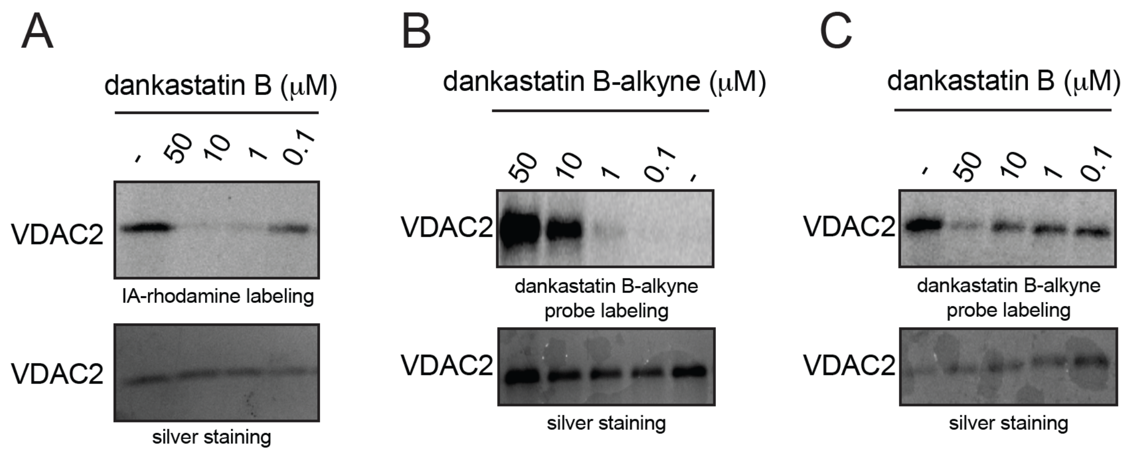


**Figure S2. Dankastatin B interactions with VDAC2. (A)** Gel-based ABPP analysis of dankastatin B against pure VDAC2 protein. Pure VDAC2 protein was pre-incubated with DMSO or dankastatin B for 30 min prior to labeling of protein with a rhodamine-functionalized iodoacetamide (IA-rhodamine) probe (1 μM) for 1 h. Proteins were resolved on SDS/PAGE and subsequently visualized by in-gel fluorescence and protein loading was assessed by silver staining. **(B)** Dankastatin B-alkyne probe labeling of VDAC2. Pure VDAC2 protein was incubated with dankastatin B-alkyne for 30 min. An azide-functionalized rhodamine was subsequently appended onto probe-labeled proteins by copper-catalyzed azide-alkyne cycloaddition, proteins were resolved by SDS/PAGE, and visualized by in-gel fluorescence. Protein loading was assessed by silver staining. **(C)** Dankastatin B-alkyne probe labeling of VDAC2 and competition of probe labeling by dankastatin B. Pure VDAC2 protein was pre-incubated with DMSO or dankastatin B for 30 min prior to incubation with dankastatin B-alkyne for 30 min. An azide-functionalized rhodamine was subsequently appended onto probe-labeled proteins by copper-catalyzed azide-alkyne cycloaddition, proteins were resolved by SDS/PAGE, and visualized by in-gel fluorescence. Protein loading was assessed by silver staining. Shown in **(B, C)** are representative gels from n=3 biological replicates/group.

### Supporting Synthetic Methods

#### General Procedures

Unless otherwise stated, all reactions were performed in oven-dried or flame-dried Fisherbrand® borosilicate glass tubes or round bottom flasks under an atmosphere of dry nitrogen. Anhydrous tetrahydrofuran, dichloromethane, *N,N*-Dimethylformamide, toluene, acetonitrile, triethylamine, and diethyl ether were obtained by passing these previously degassed solvents through activated alumina columns. All other solvents were used directly from Sigma-Aldrich Sure/Seal™ bottles. 2-Methyloct-7-yn-1-ol and **SI-7** were prepared according to literature proceedures.[1,2] Reactions were monitored by thin layer chromatography (TLC) on TLC silica gel 60 $F_{254}$ glass plates (EMD Millipore) and visualized by UV irradiation and staining with *p*-anisaldehyde, phosphomolybdic acid, or potassium permanganate. Volatile solvents were removed under reduced pressure using a rotary evaporator. Flash column chromatography was performed using Silicycle F60 silica gel (60Å, 230-400 mesh, 40-63 μm). Ethyl acetate and hexanes were purchased from Fisher Chemical and used for chromatography without further purification. Proton nuclear magnetic resonance ($^1$H NMR) and carbon nuclear magnetic resonance ($^{13}$C NMR) spectra were recorded on a Bruker AV-600 spectrometer operating at 600 MHz for $^1$H, and 150 MHz for $^{13}$C. Chemical shifts are reported in parts per million (ppm) with respect to the residual solvent signal $CDCl_3$ ($^1$H NMR: δ = 7.26; $^{13}$C NMR: δ = 77.16). High-resolution mass spectra (HRMS) were obtained by the QB3/chemistry mass spectrometry facility at the University of California, Berkeley.

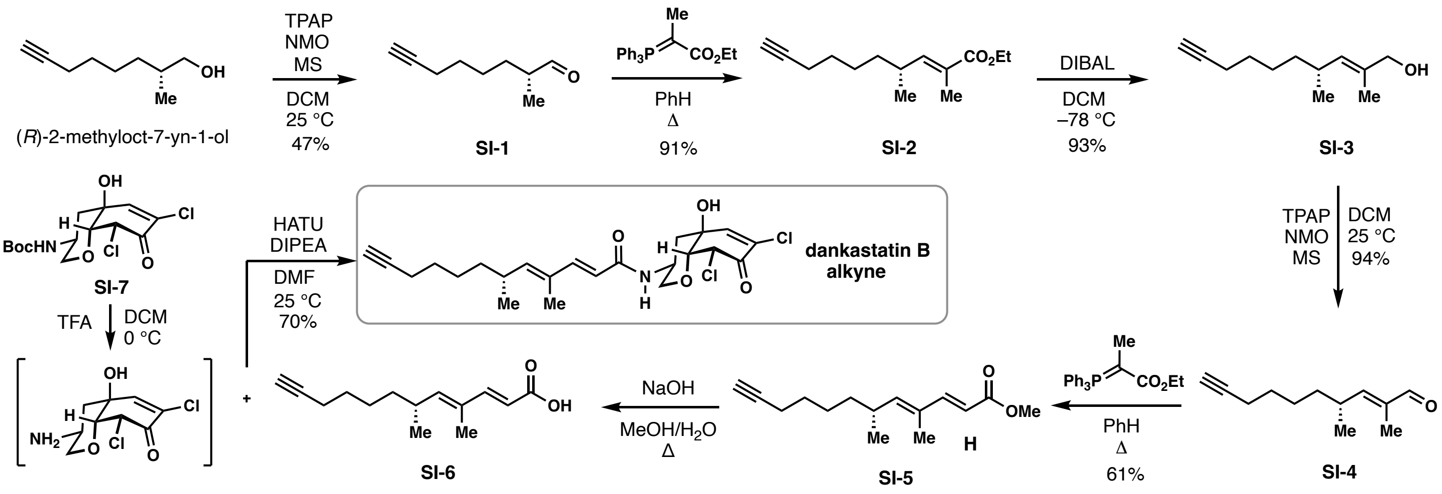


**Figure S3. Synthetic scheme for dankastatin B alkyne probe.**

**Compound SI-1.** *(R)*-2-methyloct-7-yn-1-ol (250 mg, 1.78 mmol) was dissolved in 15 mL of anhydrous DCM and to this solution was added Tetrapropylammonium perruthenate (7.3 mg, 18 μmol), 4-methylmorpholine 4-oxide (632 mg, 5.35 mmol), and powdered molecular sieves (500 mg). The reaction mixture was stirred at room temperature for 8 hours under an atmosphere of argon. Upon reaction completion as indicated by TLC, the reaction mixture was filtered through a pad of $SiO_2$ and the resulting solution concentrated *in vacuo*. The residue was purified by column chromatography (0 to 60% $Et_2O$/hexanes) to afford aldehyde **SI-1** as a pale-yellow oil (115 mg, 47%): $R_f$ = 0.72 (30% EtOAc/hexanes); IR (thin film) $v_{max}$ = 3294, 2936, 2862, 2714, 1724, 1461, 1115, 1078, 634 $cm^{-1}$; $^1H$ NMR (600 MHz, $CDCl_3$) δ 9.63 (d, *J* = 1.9 Hz, 1H), 2.35 (hd, *J* = 6.9, 2.0 Hz, 1H), 2.21 (td, *J* = 7.0, 2.7 Hz, 2H), 1.94 (t, *J* = 2.7 Hz, 1H), 1.79 – 1.66 (m, 1H), 1.61 – 1.34 (m, 5H), 1.11 (d, *J* = 7.0 Hz, 3H); $^{13}C$ NMR (151 MHz, $CDCl_3$) δ 205.0, 84.1, 77.2, 77.0, 76.7, 68.4, 46.1, 29.9, 28.3, 25.9, 18.2, 13.3; HRMS (*m/z*): (ESI) calcd. for $C_9H_{13}O$ $[M–H]^+$: 137.0966, found 137.0965.

**Compound SI-2**. Aldehyde **SI-1** (102 mg, 0.74 mmol) was dissolved in 6 mL of anhydrous benzene and (Carbethoxyethylidene)triphenylphosphorane (802 mg, 2.21 mmol) was added in one portion. The reaction mixture was heated to reflux and maintained at this temperature for 15 hours. Upon completion, as indicated by TLC, the reaction mixture was cooled to room temperature and directly purified by silica gel column chromatography (0 to 40% $Et_2O$/hexanes) to afford ester **SI-2** (150 mg, 91%): $R_f$ = 0.75 (15% $Et_2O$/hexanes); IR (thin film) $v_{max}$ = 2934, 2862, 1700, 1649, 1458, 1243, 1209, 1176, 751, 632 $cm^{-1}$; $^1H$ NMR (600 MHz, $CDCl_3$) δ 6.53 (dd, *J* = 10.1, 1.6 Hz, 1H), 4.19 (q, *J* = 7.1 Hz, 2H), 2.54 – 2.44 (m, 1H), 2.18 (td, *J* = 7.1, 2.6 Hz, 2H), 1.93

(t, $J$ = 2.6 Hz, 1H), 1.84 (s, 3H), 1.55 – 1.47 (m, 2H), 1.44 – 1.25 (m, 7H), 1.00 (d, $J$ = 6.6 Hz, 3H); $^{13}C$ NMR (151 MHz, $CDCl_3$) δ 168.8, 148.1, 126.9, 84.8, 77.6, 77.4, 77.2, 68.6, 60.8, 36.6, 33.5, 28.9, 27.0, 20.4, 18.7, 14.7, 12.9; HRMS (*m/z*): (ESI) calcd. for $C_{14}H_{23}O_2$ $[M+H]^+$: 223.1693, found 223.1693.

**Compound SI-3**. Ester **SI-2** (150 mg, 0.68 mmol) was dissolved in anhydrous DCM and cooled to –78 °C under argon. A solution of DIBAL (1M in hexane, 1.48 mL, 1.48 mmol) was added dropwise and the reaction mixture stirred at –78 °C. Upon completion, as indicated by TLC, the reaction mixture was quenched with saturated aqueous Rochelle's salt solution, warmed to room temperature, and vigorously stirred for 2 hours at room temperature. The aqueous layer was extracted with $Et_2O$ (3 x 50 mL) and the combined organic layers dried over $Na_2SO_4$ and concentrated *in vacuo*. The residue was purified by column chromatography (0 to 30% EtOAc/hexanes) to afford alcohol **SI-3** as an oil (120 mg, 98%): $R_f$ = 0.25 (15% EtOAc/hexanes); IR (thin film) $\nu_{max}$ = 3307, 2929, 2861, 1455, 1376, 1068 $cm^{-1}$; $^1H$ NMR (600 MHz, $CDCl_3$) δ 5.17 (dp, $J$ = 9.5, 1.4 Hz, 1H), 3.99 (dd, $J$ = 6.0, 1.2 Hz, 2H), 2.38 (dtdd, $J$ = 13.4, 6.7, 3.0, 1.3 Hz, 1H), 2.21 – 2.13 (m, 2H), 1.93 (t, $J$ = 2.7 Hz, 1H), 1.67 (d, $J$ = 1.4 Hz, 3H), 1.55 – 1.45 (m, 2H), 1.41 – 1.17 (m, 5H), 0.94 (dd, $J$ = 6.7, 2.8 Hz, 3H); $^{13}C$ NMR (151 MHz, $CDCl_3$) δ 133.3, 132.6, 84.6, 69.0, 68.0, 36.9, 31.9, 28.6, 26.5, 20.9, 18.3, 13.8; HRMS (*m/z*): (ESI) calcd. for $C_{12}H_{18}$ $[M–H_2O]^+$: 162.1409, found 162.1406.

**Compound SI-4**. Alcohol **SI-3** (110 mg, 0.61 mmol) was dissolved in 6 mL of anhydrous dichloromethane followed by the addition of Tetrapropylammonium perruthenate (2.4 mg, 6.1 μmol), 4-methylmorpholine 4-oxide (216 mg, 1.83 mmol) and powdered molecular sieves (500 mg). The reaction mixture was stirred at room temperature until starting material was consumed, as indicated by TLC. The reaction mixture was filtered through a pad of celite and the filtrate rinsed with DCM. The solvent was removed *in vacuo* and the residue purified by column chromatography (0 to 20% EtOAc/hexanes) to give aldehyde **SI-4** as an oil (102 mg, 94%): $R_f$ = 0.72 (40% EtOAc/hexanes); IR (thin film) $\nu_{max}$ = 2929, 2858, 1730, 1688, 1642, 1459, 1324, 1017, 632 $cm^{-1}$; $^1H$ NMR (600 MHz, $CDCl_3$) δ 9.39 (s, 1H), 6.25 (dq, $J$ = 9.9, 1.4 Hz, 1H), 2.75 – 2.65 (m, 1H), 2.18 (td, $J$ = 7.0, 2.6 Hz, 2H), 1.93 (t, $J$ = 2.6 Hz, 1H), 1.75 (d, $J$ = 1.4 Hz, 3H), 1.55 – 1.33 (m, 6H), 1.07 (d, $J$ = 6.7 Hz, 3H); $^{13}C$ NMR (151 MHz, $CDCl_3$) δ 195.5, 160.1, 138.1, 84.2, 77.2, 77.0, 76.8, 68.4, 36.1, 33.4, 28.4, 26.4, 19.8, 18.2, 9.4; HRMS (*m/z*): (ESI) calcd. for $C_{12}H_{17}O$ $[M–H]^+$ : 177.1276, found 177.1277.

**Compound SI-5**. Aldehyde **SI-4** (97 mg, 0.54 mmol) was dissolved in 6 mL of anhydrous toluene. To this solution was added Methyl (triphenylphosphoranylidene)acetate (565 mg, 1.63 mmol) and the resulting solution was heated at 90 °C. Upon completion, as indicated by TLC, the reaction mixture was cooled to room temperature and directly purified by silica gel chromatography (20% $Et_2O$/hexanes) to afford ester **SI-5** as an oil (76 mg, 60%): $R_f$ = 0.51 (15% $Et_2O$/hexanes); IR (thin film) $v_{max}$ = 2932, 2861,1717, 1623, 1455, 1434, 1394, 1270, 1169, 847 $cm^{-1}$; $^1H$ NMR (600 MHz, $CDCl_3$) δ 7.31 (d, *J* = 15.7 Hz, 1H), 5.79 (d, *J* = 15.7 Hz, 1H), 5.67 (d, *J* = 9.8 Hz, 1H), 3.75 (s, 3H), 2.58 – 2.48 (m, 1H), 2.17 (tdd, *J* = 7.1, 2.7, 1.1 Hz, 2H), 1.93 (q, *J* = 2.8 Hz, 1H), 1.78 (s, 3H), 1.57 – 1.22 (m, 6H), 0.99 (d, *J* = 6.6 Hz, 3H); $^{13}C$ NMR (151 MHz, $CDCl_3$) δ 168.2, 150.3, 148.5, 131.6, 115.4, 84.6, 77.4, 77.2, 77.0, 68.4, 51.6, 36.8, 33.3, 28.7, 26.7, 20.6, 18.5, 12.5; HRMS (*m/z*): (ESI) calcd. for $C_{15}H_{23}O$ $[M+H]^+$: 235.1693, found 235.1694.

**Compound SI-6**. To ester **SI-5** (75.0 mg, 0.32 mmol) was added 4.8 mL of MeOH and 1.0 mL of $H_2O$. To this suspension was added NaOH (64.0 mg, 1.60 mmol) and the reaction mixture stirred at 35 °C. Upon reaction completion, as indicated by TLC, the reaction mixture was cooled to 0 °C and 1N HCl (10 mL) and EtOAc (10 ml) were added. The aqueous layer was extracted with ethyl acetate (3 x 25 mL) and the combined organic layers were combined, dried over $Na_2SO_4$ and concentrated *in vacuo*. The resulting residue was purified by column chromatography (30% to 100% EtOAc/hexanes) to afford acid **SI-6** as a pale-yellow oil (65 mg, 92%): $R_f$ = 0.56 (3.5% MeOH/Dichloromethane); IR (thin film) $v_{max}$ = 3305, 2935, 2861, 2684, 2593, 1685, 1617, 1208, 851 $cm^{-1}$; $^1H$ NMR (600 MHz, $CDCl_3$) δ 7.39 (d, *J* = 15.6 Hz, 1H), 5.79 (d, *J* = 15.6 Hz, 1H), 5.72 (d, *J* = 9.8 Hz, 1H), 2.60 – 2.50 (m, 1H), 2.17 (tdd, *J* = 7.1, 2.7, 1.1 Hz, 2H), 1.93 (t, *J* = 2.6 Hz, 1H), 1.80 (d, *J* = 1.2 Hz, 3H), 1.51 (dq, *J* = 8.7, 6.3 Hz, 2H), 1.44 – 1.24 (m, 4H), 1.00 (d, *J* = 6.7 Hz, 3H); $^{13}C$ NMR (151 MHz, $CDCl_3$) δ 172.9, 152.4, 149.8, 131.7, 114.9, 84.6, 68.4, 36.7, 33.4, 28.7, 26.7, 20.5, 18.5, 12.5; HRMS (*m/z*): (ESI) calcd. for $C_{14}H_{21}O_2$ $[M+H]^+$: 221.1536, found 221.1536.

**Dankastatin B alkyne**. Alcohol **SI-7** (3 mg, 8.5 µmol) was dissolved in anhydrous DCM (0.5 mL) and the resulting solution was cooled to 0 °C wherein 50 µL of trifluoracetic acid was added. The reaction mixture was stirred at 0 °C for an hour at which point additional TFA (50 µL) was added and the reaction warmed to room temperature and stirred for 30 additional minutes. The volatiles were removed *in vacuo* to afford a yellow oil which was redissolved in 0.7 mL of anhydrous DMF. To this solution was added acid **8** (2.1 mg, 9.3 µmmol), DIPEA (5.9 µL, 35.7 µmol), HATU (3.6 mg, 9.4 µmol) and the reaction mixture was stirred at room temperature. Upon completion, as indicated by TLC, the reaction mixture was diluted with 5 mL of $Et_2O$ and quenched with saturated ammonium chloride solution (10 mL). The aqueous layer was extracted with $Et_2O$ (3 x 10 mL) and the combined organic layers were dried over $Na_2SO_4$ and concentrated *in vacuo*. The resulting residue was purified by preparative thin layer chromatography (35% Acetone/hexanes) to afford **Dankastatin B alkyne** as a film (2.7 mg, 70%): $R_f$ = 0.46 (35% Acetone/hexanes); IR (thin film) $v_{max}$ = 3379, 3295, 2932, 2856, 1721, 1649, 1609, 1528, 1107, 748 $cm^{-1}$; $^1H$ NMR (700 MHz, $CDCl_3$) δ 7.24 (d, *J* = 15.3 Hz, 1H), 6.82 (d, *J* = 2.5 Hz, 1H), 5.79 – 5.66 (m, 2H), 5.32 (d, *J* = 2.2 Hz, 1H), 5.25 (d, *J* = 8.0 Hz, 1H), 4.20 – 4.10 (m, 2H), 3.97 (t, *J* = 2.4 Hz, 1H), 3.24 (t, *J* = 10.8 Hz, 1H), 2.88 (s, 1H), 2.57 – 2.50 (m, 2H), 2.19 (tt, *J* = 7.1, 2.4 Hz, 2H), 1.96 (t, *J* = 2.6 Hz, 1H), 1.82 – 1.74 (m, 4H), 1.56 – 1.49 (m, 2H), 1.45 – 1.33 (m, 2H), 1.35 – 1.26 (m, 2H), 1.01 (dd, *J* = 6.7, 2.1 Hz, 3H); $^{13}C$ NMR (151 MHz, $CDCl_3$) δ 182.9, 166.6, 148.5, 147.8, 142.4, 133.9, 131.1, 116.9, 84.6, 83.9, 70.1, 69.9, 68.4, 61.4, 44.2, 44.1, 36.8, 33.3, 28.7, 26.8, 20.6, 18.5, 12.7; HRMS (*m/z*): (ESI) calcd. for $C_{23}H_{30}O_4NCl_2$ $[M+H]^+$: 454.1546, found 454.1542.

**Supporting Information References**


(1) Siow, A.; Opiyo, G.; Kavianinia, I.; Li, F. F.; Furkert, D. P.; Harris, P. W. R.; Brimble, M. A. Total Synthesis of the Highly *N*-Methylated Acetylene-Containing Anticancer Peptide Jahanyne. *Org. Lett*. **2018**, *20*, 788.

(2) Tong, B.; Belcher, B. P.; Nomura, D. K.; Maimone, T. J. Chemical investigations into the biosynthesis of the gymnastatin and dankastatin alkaloids. *Chem. Sci*. **2021**, *12*, 8884.
